# MYST acetyltransferases are a targetable therapeutic vulnerability in SETBP1-mutant leukemia

**DOI:** 10.64898/2026.01.08.697228

**Authors:** Hanqian L Carlson, Thai T Nguyen, Samantha Tauchmann, Sarah A Carratt, Traci L Kruer, Nana Adjoa Ben-Crentsil, Maria Balasis, Hyeyoon Kim, Chia-Feng Tsai, Tao Liu, Shawn B Shrestha, Jared M Fischer, Eric Padron, Theodore P Braun, Julia E Maxson

**Author notes:** Correspondence to: -Dr. Julia Maxson, Oregon Health & Science University Mail Code: KR-HEM, 3181 SW Sam Jackson Park Road, Portland, OR 97239. -Dr. Theodore Braun, Oregon Health & Science University Mail Code: KR-HEM, 3181 SW Sam Jackson Park Road, Portland, OR 97239.

## Abstract

Mutations in SET binding protein 1 (*SETBP1*) are associated with an adverse prognosis in myeloid malignancies. These mutations stabilize SETBP1 protein, driving increased expression of a progenitor-associated gene expression program through incompletely described mechanisms. A proteomic screen revealed interactions between SETBP1 and members of the MYST acetyltransferase complexes, including the catalytic subunits—KAT6A and KAT7. Mutant SETBP1 increases the localization of MYST complexes at known SETBP1 target genes, including the *Hoxa* cluster, where it drives increased histone acetylation and gene expression. Treatment of *SETBP1^D868N^*-expressing myeloid progenitors with MYST inhibitors reduced target gene expression. To establish the efficacy of MYST inhibition *in vivo*, we treated mice harboring a syngeneic *SETBP1*-mutant leukemia with the clinical-grade MYST inhibitor—PF-9363. This resulted in complete hematologic control and increased survival. MYST inhibition was also highly effective against a *SETBP1*-mutant PDX model. These studies identify MYST acetyltransferases as promising therapeutic targets in *SETBP1*-mutant malignancies.

**Statement of Significance:** SETBP1 mutations are markers of high-risk myeloid malignancies, but we lack any targeted therapies to improve outcomes. In this study, we identify MYST acetyltransferases as key drivers of mutant SETBP1-driven transcription. MYST inhibitors are highly effective against SETBP1-mutant leukemia and represent a promising avenue for clinical translation.

## Introduction

SETBP1 is a transcriptional regulatory protein that is mutated in myeloid malignancies. This includes 45% of Chronic Neutrophilic Leukemia (CNL) cases, 33% of myelodysplastic/myeloproliferative neoplasms with neutrophilia (MDS/MPN-N), 4-15% of chronic myelomonocytic leukemia (CMML), and 30% of juvenile myelomonocytic leukemia^1–7^. Mutations in *SETBP1* are often associated with adverse survival in these contexts^4,6^. Additionally, *SETBP1* mutations are associated with the emergence of chemotherapy-resistant clones in Juvenile Myelomonocytic Leukemia, decreasing the 5-year event-free survival of children with this disease from 51% to 18%^4^. In addition to these myeloproliferative disorders, *SETBP1* mutations are also found in acute myeloid leukemia (AML). Consistent with an association with more aggressive disease, *SETBP1* mutations are found at low frequency (∼1%) in *de novo* AML^8,9^, but are frequent in secondary AML that evolved from a prior myeloid malignancy (∼17% of cases)^5^. Collectively, these data suggest that *SETBP1* mutations are prevalent in myeloid malignancies and are associated with extremely poor patient outcomes.

Mutations in *SETBP1* occur in a restricted region of the protein associated with a recognition motif for the E3-ubiquitin ligase, β-TrCP1^3^. Mutations in this motif are thought to prevent ligase recognition, thus reducing ubiquitination and protein turnover^3^. The resulting stabilization of SETBP1 protein drives its oncogenic effects. In solid tumors, SETBP1 has been implicated as a negative regulator of the tumor suppressor protein phosphatase 2A (PP2A). SETBP1 was originally identified in a yeast two-hybrid screen as an interacting partner of its namesake—S/E Translocation (SET), and they cooperate to induce degradation of the tumor suppressor PP2A^3,10^. Degradation of PP2A, an important cell cycle regulatory protein, enhances cell division. Although this is likely an important function of SETBP1 in some tumor contexts, deletion of the SET-interacting domain does not disrupt the oncogenic potential of SETBP1 in myeloproliferative models^11^, indicating that additional SETBP1 functions exist.

In blood cancers, SETBP1 mutations have been associated with pro-leukemogenic transcriptional changes. This includes increased expression of transcriptional regulators *Hoxa9*, *Hoxa10*, *Myc*, *Myb*, *Meis1*, and *Mecom*^11–15^. These transcriptional regulators are classically associated with stem and progenitor gene expression programs, and their overexpression drives leukemic transformation in other contexts. The mechanism by which SETBP1 controls transcription is an area of active investigation. SETBP1 has three AT-hook DNA-binding motifs. Two recent studies identified an interaction between SETBP1 and the leukemia-associated chromatin regulatory protein MLL1 (aka KMT2A), a histone methyltransferase^15,16^. MLL fusion proteins are associated with leukemia development and also lead to increased expression of *HoxA9*, *HoxA10, and Meis1*^17^. MLL1 was found to be important for the transcription of these genes in *SETBP1-*mutant cells^16^. Although MLL1 clearly plays an important role, whether other chromatin proteins are required for *SETBP1*-driven transcriptional changes is unknown.

In the present work, we use a proteomics-based approach to map the SETBP1-interactome, which revealed robust interaction between SETBP1 and MYST acetyltransferase complexes, which deposit activating histone marks. Here we report that mutant SETBP1 increases the localization of MYST acetyltransferase complexes at stem and progenitor-associated genes and increases deposition of gene-activating histone acetyl marks. Furthermore, we find that the MYST acetyltransferases KAT6A and KAT7 are key dependencies in SETBP1-mutant cells and that clinically relevant MYST inhibitors are potent against SETBP1-mutant leukemia using a combination of murine and patient-derived xenograft models. These studies detail an intricate mechanism of SETBP1-driven gene regulation and nominate KAT6A and KAT7 as promising therapeutic targets in *SETBP1*-mutant disease.

## RESULTS

### A proteomic screen identifies transcriptional regulatory complexes that interact with SETBP1

To understand SETBP1’s role in gene regulation in greater depth, we employed biotin-labelling identification (BioID) to map the SETBP1 interactome^18–21^. In this approach, “bait” protein is expressed and fused to the *E. coli* biotin ligase, BirA. BirA biotinylates nearby proteins within a labelling radius of ∼20 nm, capturing both stable and transient interactions^20^. We generated SETBP1^WT^ and SETBP1^D868N^–BirA* constructs and introduced them into a human hematopoietic cell line (K562). Biotinylated proteins were purified on streptavidin beads and identified using high-sensitivity mass spectrometry. The SETBP1^WT^-BirA, SETBP1^D868N^-BirA, and BirA control conditions were performed in quadruplicate with strong concordance between replicates (Figures 1 and Supplemental Table 1).

**Figure 1.**
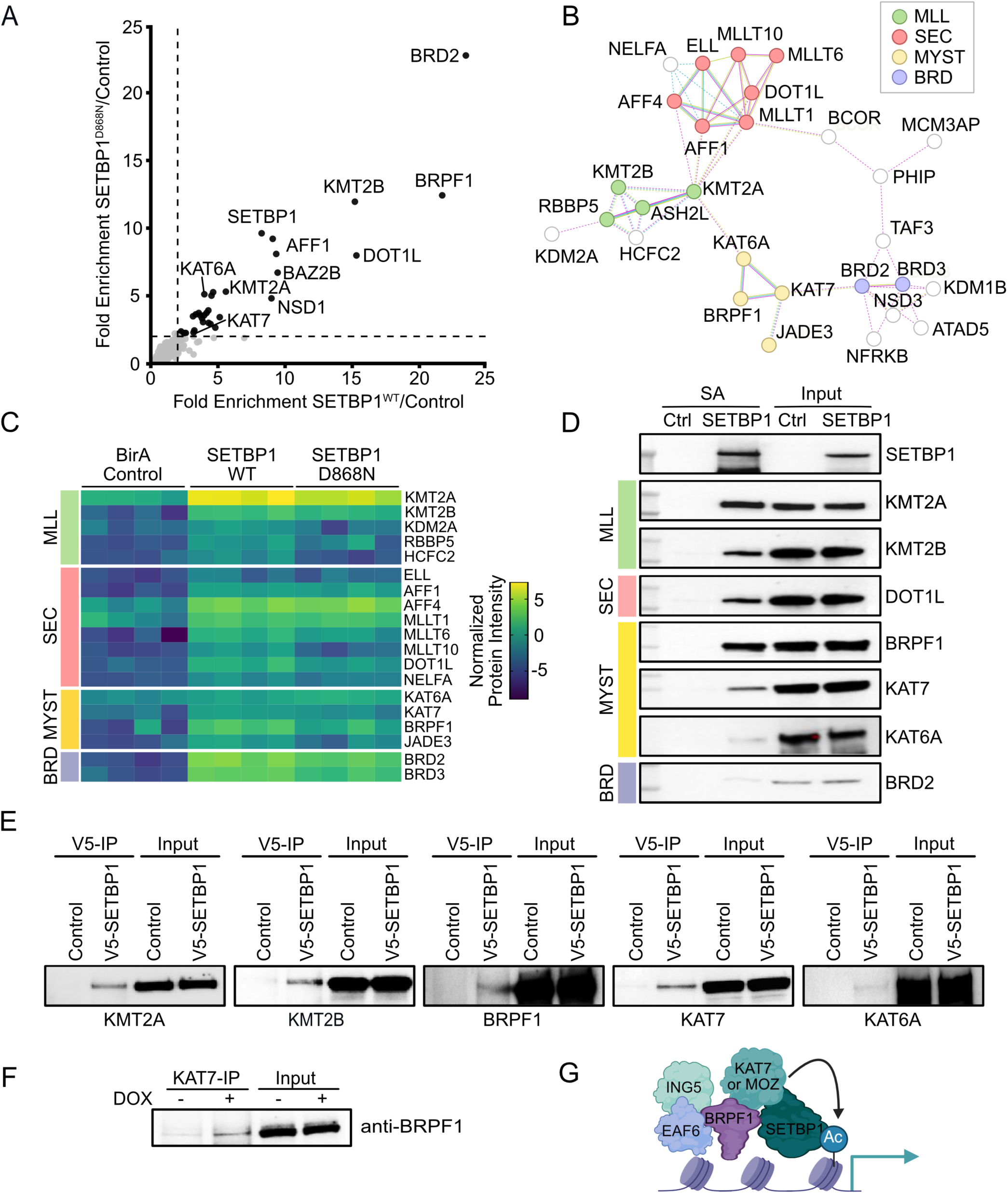
BioID-based screen identifies epigenetic regulatory complexes in close proximity to SETBP1. **A.** Plot of fold enrichment of mean biotinylated protein abundance derived from four replicates per condition. Plot represents the enrichment of peptides from control vector-BirA*, *SETBP1^WT^*-BirA*, and *SETBP1^D868N^*-BirA*. The x-axis of the presented plot shows the fold enrichment in SETBP1^WT^ versus control. The y-axis of the plot shows fold enrichment in SETBP1^D868N^ versus control. Dashed lines represent a fold enrichment value of 2. **B.** Network of proteins identified in *SETBP1^WT^*-BirA* or *SETBP1^D868N^*-BirA* versus Control-BirA* transduced K562 cells. Proteins with >2-fold enrichment over BirA alone in either of the *SETBP1* conditions were analyzed using STRING, a physical interaction network. Clusters were identified using DBScan, and enriched terms within a cluster are indicated by the labels. The color of the nodes represents the identified pathways to which the proteins belong to as described in the figure. Proteins involved in chromatin biology are highlighted by color—green: histone methyltransferase, pink: super elongation complex, yellow: histone acetyltransferase, blue: bromodomain proteins. **C.** Heatmap of normalized intensity for key proteins identified in BioID in *SETBP1^WT^*-BirA* or *SETBP1^D868N^*-BirA* versus Control-BirA* transduced K562 cells as prioritized in the network shown in (B). Four replicates are shown for each condition. **D.** Validation of hits from the screen by streptavidin (SA) pulldown followed by immunoblot with the indicated antibodies in K562 cells expressing *SETBP1^WT^*-BirA* or a BirA* control. **E**. Co-immunoprecipitation of V5-tagged control or V5-tagged-SETBP1 in HEK293 cells. The V5 antibody was used for immunoprecipitation, followed by immunoblots with antibodies against the indicated putative SETBP1-interaction partners. **F.** Co-immunoprecipitation of BRPF1 and KAT7 in control (-DOX) and SETBP1^D868N^-induced (+ DOX) conditions in K562 cells demonstrates enhanced interaction of KAT7 and BRPF1 in *SETBP1*^D868N^-expressing cells**. G.** Proposed model of MYST complex interaction with SETBP1.

To interrogate this dataset, we plotted the fold enrichment of normalized counts for SETBP1^WT^ and SETBP1^D868N^ relative to the control. Of the 1209 proteins quantified in this dataset, 37 proteins had a 2-fold or greater enrichment for both SETBP1^WT^ and SETBP1^D868N^, as compared to the control condition (Figure 1A). High concordance between the relative fold enrichment indicates that wildtype and mutant SETBP1 have broadly similar interactomes (Figure 1A), consistent with the idea that mutations in SETBP1 act by increasing SETBP1 abundance. Identification of a known SETBP1-interacting protein, KMT2A (aka MLL1), provided confidence in the robustness of this dataset^15,16^.

This screen identified a number of epigenetic regulatory proteins in close proximity to SETBP1. To understand the relationships between putative SETBP1-interacting proteins, proteins with a significant enrichment over BirA-Control in either of the *SETBP1* conditions were analyzed with STRING using a physical interaction network^22^. Clusters were identified using DBScan, and enriched terms within a cluster are indicated by the labels (Figure 1B). This clustering identified putative interactions between SETBP1 and the following transcriptional regulatory complexes: 1) histone methyltransferase MLL, 2) super elongation complex, 3) histone acetyltransferase MYST, and 4) bromodomain proteins BRD. A heatmap of normalized intensity illustrates that SETBP1^WT^ and SETBP1^D868N^ have similar key interaction partners (Figure 1C). We validated candidate interaction partners from each of these 4 protein complexes by BioID-streptavidin pull-down followed by Western blotting (Figure 1D).

The two most enriched proteins identified in this screen were the bromodomain protein BRD2 and the MYST complex member BRPF1 (Figure 1A, C). In prior studies, however, we found that inhibitors of BRD2 do not impair the growth of *SETBP1*-mutant cells^11^. We therefore focused our attention on BRPF1 and the MYST acetyltransferase complex. We also identified two MYST catalytic subunits: KAT7 (aka HBO1) and KAT6A (aka MOZ) (Figure 1A-D). Co-immunoprecipitation studies validated a robust association of SETBP1 with BRPF1, KAT6A, and KAT7 (Figure 1E). Because *SETBP1^D868N^* binds to both BRPF1 and KAT7, we postulated that it might enhance their interaction to promote complex activity. A co-immunoprecipitation experiment revealed increased pulldown of BRPF1 by KAT7 in K562 cells expressing *SETBP1^D868N^* (Figure 1F). These studies provide the first comprehensive map of the SETBP1 interactome and nominate MYST acetyltransferases as candidate mediators of SETBP1-driven transcriptional effects (Figure 1G).

### SETBP1 and MYST co-localize on chromatin

Having identified a robust interaction between SETBP1 and MYST complexes we next set out to understand whether they are bound at overlapping genomic regions. We first performed CUT&RUN for SETBP1^D868N^ in an inducible K562 cell model. Induction resulted in a robust detection of chromatin-bound SETBP1 (Figure 2A). SETBP1 has three AT-hook motifs that canonically bind to the minor groove of DNA in AT-rich regions. Consistent with this, we find that 96.29% of SETBP1-bound regions contain an AT-rich motif (Figure 2B).

**Figure 2.**
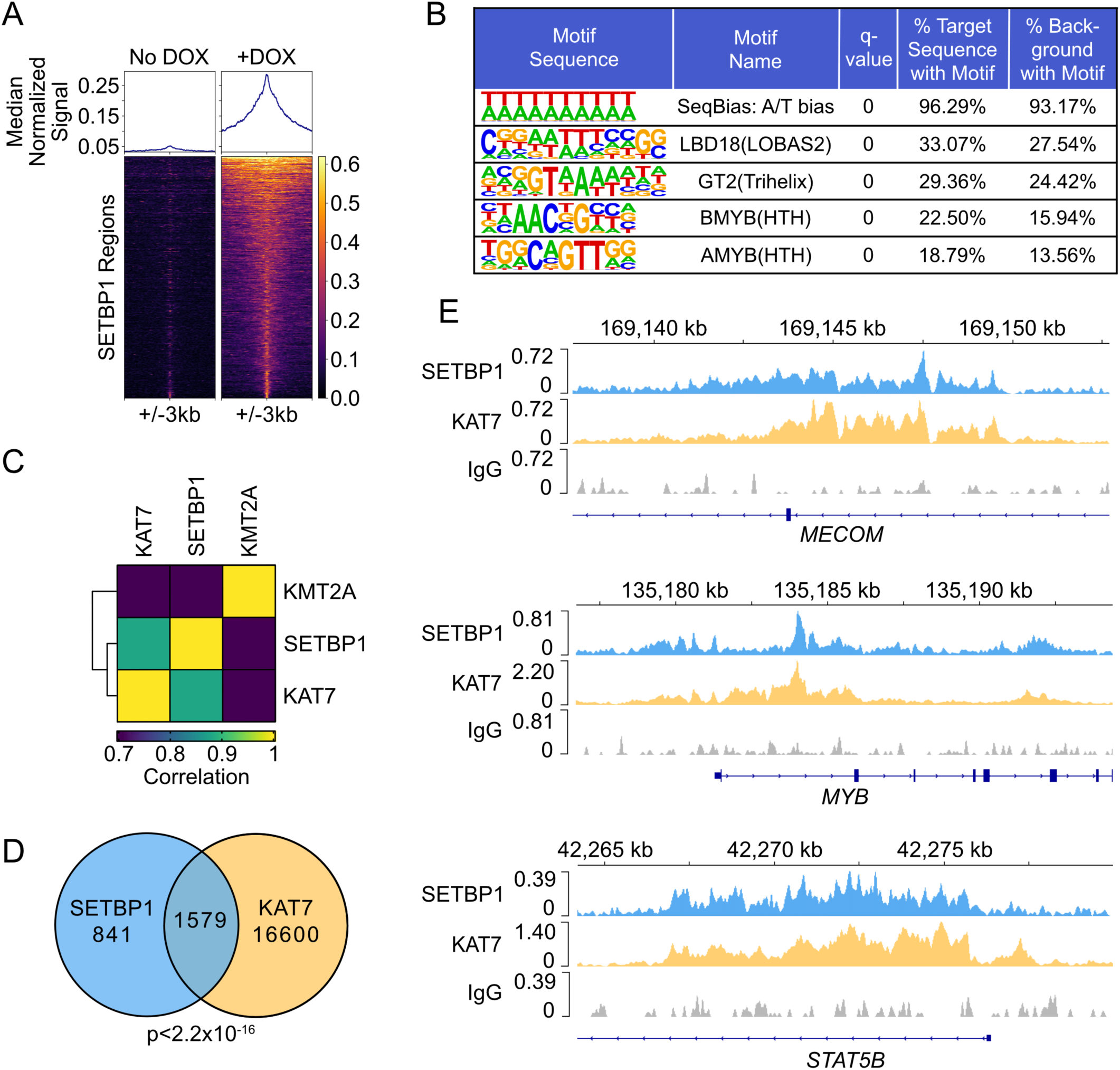
The majority of SETBP1 binding sites are also bound by the MYST catalytic subunit, KAT7. **A.** CUT&RUN for SETBP1 in K562 cells with a doxycycline (DOX)-inducible construct in triplicate. Heatmaps and median signal profiles of normalized SETBP1 CUT&RUN signal in control (no DOX) and SETBP1^D868N^-induced (+DOX) conditions. The signal is shown at peak centers +/- 3kb. Each row in the heatmap represents one SETBP1 binding region. **B.** Top motifs identified by HOMER at SETBP1-bound regions. The q-values represent Benjamini-Hochberg corrected p-values calculated by HOMER. Percentages indicate the fraction of sequences in target or background regions that contain each motif. Motifs with q-value<0.0001 were ranked in descending order by percentage of target regions (SETBP1 peaks) containing the motif, and the top five are shown. **C.** Spearman correlation of SETBP1, KAT7, and KMT2A signal in SETBP1^D868N^-induced condition. SETBP1 has a high degree of correlation with KAT7. **D.** Venn diagram of SETBP1 and KAT7 binding regions in the SETBP1^D868N^-induced condition. There are 841 regions bound by SETBP1 only, 16600 regions bound by KAT7 only, and 1579 regions co-occupied by both SETBP1 and KAT7. E. Normalized signal tracks for SETBP1, KAT7, and IgG control at *MECOM*, *MYB*, and *STAT5B* loci in the SETBP1^D868N^-induced condition.

To understand whether SETBP1^D868N^ and MYST complexes co-localize on chromatin, we further mapped the binding of a representative MYST catalytic subunit (KAT7). Genome-wide localization of a known SETBP1-interaction partner, KMT2A^15,16^, serves as a positive control. We find a high degree of correlation between SETBP1 ^D868N^-bound regions and both KMT2A and KAT7-bound regions (Figure 2C). Of the 2,420 SETBP1^D868N^-bound regions, approximately two-thirds of these (1,579 regions) overlapped with KAT7 binding—a highly significant overlap (p<2.2e-16, Figure 2D). Genomic feature analysis revealed that SETBP1 and KAT7 colocalized primarily at promoters (55%) and to a lesser extent at intergenic regions (Figure S1A). Examples of overlapping peaks for SETBP1 and KAT7 are shown at the *MYB*, *MECOM*, and *STAT5B* genes (Figure 2E). Taken together, these studies demonstrate that the majority of SETBP1 peaks across the genome are co-bound by KAT7.

### SETBP1^D868N^ upregulates proliferation and stem-associated gene expression programs in murine hematopoietic progenitors

Although K562 cells are a highly tractable model for biochemistry, we next sought a primary hematopoietic cell model that would allow us to evaluate the effects of SETBP1-associated epigenetic complexes on transcriptional control. To this end, murine lineage-negative hematopoietic stem and progenitor cells were transduced with *SETBP1^WT^*, *SETBP1^D868N^*, or a *luciferase*-expressing control vector with a GFP marker (Figure 3A). A colony-forming unit (CFU) assay on sorted cells revealed that exogenous expression of either *SETBP1*^WT^ or *SETBP1*^D868N^ significantly augmented colony formation, with the most pronounced increase in the *SETBP1*^D868N^ condition (Figure 3B-C). This is consistent with mutations in SETBP1 elevating protein levels to support oncogenesis. The SETBP1-driven colonies had large, dense centers (Figure 3B). Evaluation of gene expression at day 14 revealed significant upregulation of stem and progenitor-associated SETBP1 target genes such as *Hoxa9*, *Hoxa10,* and *Mecom* in *SETBP1*-expressing cells (Figure 3D). We next evaluated transduced progenitors in a parallel liquid culture model. In liquid culture, we observed limited proliferation of the control-vector-transduced progenitors until day 7. *SETBP1^WT^* expressing progenitors had markedly increased growth over the control, but plateaued at day 11, whereas *SETBP1^D868N^* progenitors continued to expand through the duration of the assay (Figure 3E). These studies establish a tractable system for evaluating *SETBP1*-associated gene expression changes in primary hematopoietic progenitors.

**Figure 3.**
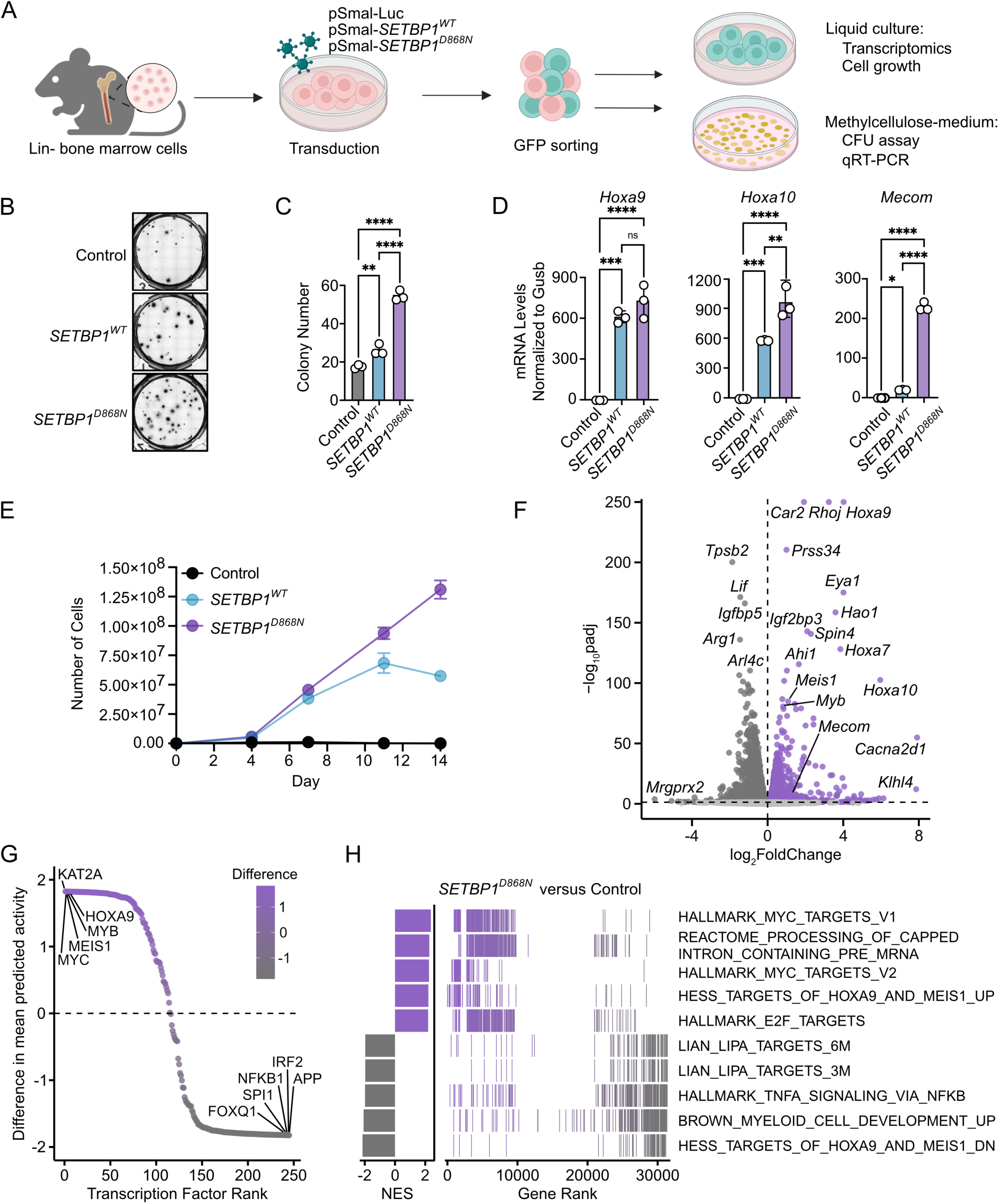
SETBP1^D868N^ upregulates proliferation and stem-associated pathways and activity in murine hematopoietic lineage-depleted cells. **A.** Schematic overview of lineage-depleted (Lin-) bone marrow isolated and transduced with a pSMAL-control-GFP lentiviral vector, pSMAL-*SETBP1*^WT^-GFP vector, or a pSMAL-*SETBP1^D868N^*-GFP vector. GFP-positive cells were sorted to enrich for transduced cells and plated in colony colony-forming unit (CFU) assay or liquid culture. **B.** These transduced cells were plated in methylcellulose-based colony-forming medium (1000 cells/well). Representative images of control, *SETBP1^WT^*, and *SETBP1^D868N^*-expressing cells are shown. **C.** The number of colonies was counted in triplicate wells after 14 days. Error bars represent SEM. Statistics: 1-way ANOVA with Dunnett’s multiple comparisons test. *p < 0.05, ** *p* < 0.01, *** *p* < 0.001, \*\*\*\**p* < 0.0001. **D.** qRT-PCR analysis of *Hoxa9*, *Hoxa10,* and *Mecom* mRNA transcripts from colonies harvested at 14 days. Expression levels were normalized to the Gusb housekeeping gene. **E.** Control, *SETBP1^WT^* or *SETBP1^D868N^* Lin-bone marrow cells were cultured in IMDM medium with 15% FBS, SCF, IL6, and IL3 for 2 weeks in triplicate wells. Proliferation of cells was determined by trypan-based automated cell counts. **F.** RNA-seq analysis of Lin-cells in *SETBP1^D868N^* and control conditions. Volcano plot of differentially expressed genes in each condition. Approximately 3100 genes were significantly upregulated (Fold change > 1.5, purple) and ∼3200 genes were downregulated (Fold change < −1.5, dark grey) in *SETBP1^D868N^* versus control cells (Padj < 0.05). Non-significant genes are in light grey. Due to numeric underflow in statistical calculations, some genes were assigned adjusted p-values of 0, resulting in infinite -log10(padj) values. For visualization, these values were capped at 250. **G.** Transcription factors (TFs) ranked by the difference in their mean predicted activity in *SETBP1^D868N^* versus control cells. Predicted activity scores were determined using Priori, and points are colored by the magnitude and direction of the difference. Top-ranked TFs with higher mean predicted activity in the *SETBP1^D868N^* condition are in purple, and TFs with higher mean predicted activity in the control (are in gray. **H.** GSEA of *SETBP1^D868N^* and control conditions using M2 and Hallmark gene sets. Left: Normalized enrichment scores (NES) for the top 5 gene sets enriched in *SETBP1^D868N^* (Purple) and control (Grey). Right: Positions of GSEA-ranked input genes within each top gene set. Colors indicate enrichment direction (purple: *SETBP1^D868N^*, grey: control).

We next set out to define the *SETBP1^D868N^*-regulated gene set in these cells to contextualize subsequent epigenetic studies. Bulk RNA-seq was used to profile gene expression changes in *SETBP1^D868N^*-transduced Lin-cells versus those transduced with the control vector, cultured for 72 hours post-sorting. Upregulated genes in *SETBP1^D868N^* expressing cells included *Hoxa9*, *Hoxa10*, *Myb*, *Meis1,* and *Mecom* (Figure 3F). Next, transcription factor activity in each condition was calculated using Priori^23^. Top-ranked candidate transcription factors predicted to be involved in the *SETBP1^D868N^*-associated gene signature were MYC, MEIS1, MYB, and HOXA9 (Figure 3G), consistent with upregulation of these hematopoietic transcription factors by SETBP1. Comparative gene ontology (GO) enrichment from *SETBP1^D868N^*-upregulated genes found the most significant Molecular Function (MF) pathways to be involved in RNA and DNA catalytic activity, protein folding chaperones, and mRNA-associated processes (Figure S1B). Gene-set enrichment analysis was performed against the KEGG database in *SETBP1^D868N^*-expressing cells versus control. The most prominent enriched gene sets were associated with MYC, HOXA9, MEIS1, and E2F signaling (Figure 3H). In contrast, differentiation-related programs were significantly down-regulated. These studies credential this murine progenitor model for the evaluation of SETBP1-driven gene regulation.

### SETBP1^D868N^ increases the deposition of MYST-associated histone marks at leukemia-associated genes

To understand the functional significance of SETBP1’s interaction with MYST complexes, we profiled the chromatin occupancy of KAT7 (a catalytic subunit of MYST) in murine progenitors expressing a control vector or *SETBP1^D868N^* using CUT&RUN. Expression of *SETBP1^D868N^* increased the binding of both KMT2A and KAT7 genome-wide (Figure 4A). In *SETBP1^D868N^*-expressing hematopoietic progenitors, increased binding of KAT7 was observed at SETBP1-regulated genes, such as the *HOXA* cluster and *MEIS1* (Figure 4B).

**Figure 4.**
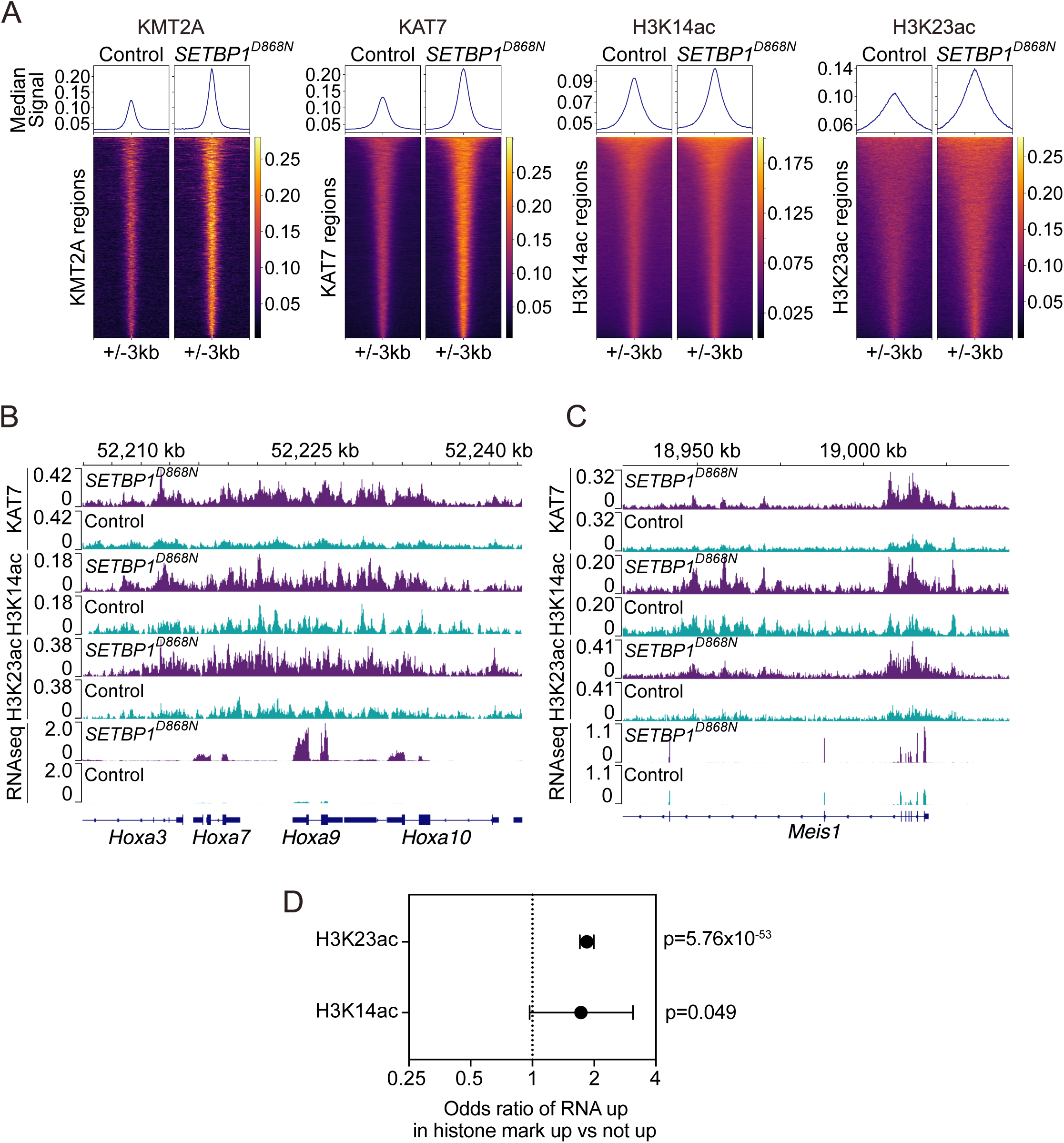
SETBP1^D868N^ increases the deposition of MYST-associated histone marks at leukemia-associated genes. Murine Lin-hematopoietic progenitors were transduced with a *SETBP1^D868N^* or control vector as described in Figure 3A and cultured for 5 days. **A.** Heatmaps and median signal profiles of normalized CUT&RUN signal for KMT2A, KAT7, H3K14ac, and H3K23ac in control and *SETBP1^D868N^* conditions. CUT&RUN signal is shown at peak centers +/- 3kb. **B.** Normalized signal tracks for KAT7, H3K14ac, H3K23ac, and RNAseq at the *Hoxa* cluster and **C.** *Meis1* locus in the *SETBP1^D868N^* and control conditions. **D.** Enrichment of genes with increased expression in RNAseq in the gene sets with increased H3K14ac/H3K23ac versus those without increased acetylation. Significance assessed by Fisher’s exact test.

MYST complexes containing KAT7/BRPF1 are primarily thought to acetylate Histone H3 Lysine 14 (H3K14ac), a marker of active gene regulatory elements^24^. KAT6A/BRPF1-containing complexes are thought to primarily acetylate Histone H4 Lysine 23 (H3K23ac)^25^. H3K14ac and H3K23ac localization were profiled by CUT&RUN in control and *SETBP1^D868N^* progenitors. Hematopoietic progenitors expressing *SETBP1^D868N^* had increased genome-wide deposition of both H3K14ac and H3K23ac (Figure 4A), including SETBP1 target genes such as the *Hoxa* cluster and *Meis1* (Figure 4B-C). Genomic feature analysis revealed that the majority of KAT7/H3K14ac/H3K23ac peaks were in promoters (Figure S2B). We next evaluated the association of this increased histone mark deposition on gene expression. Genes with increased expression by RNAseq were significantly enriched within those genes with increased H3K14ac or H3K23ac (1.73 fold and 1.84 fold, respectively) (Figure 4D). These data suggest that SETBP1^D868N^ leads to increased histone acetylation by MYST complexes to activate transcription.

### SETBP1^D868N^-driven proliferation is dependent on MYST acetyltransferase activity

We next wanted to determine whether *SETBP1^D868N^*-mutant cells depend on MYST acetyltransferases for their growth and leukemogenic transcriptional programming. We treated *SETBP1^D868N^*-expressing murine lineage-negative progenitors with PF-9363, which inhibits both KAT7 and KAT6A, or WM-3835, a specific KAT7 inhibitor. Treatment of these progenitors with PF-9363 reduced *SETBP1^WT^* or *SETBP1^D868N^*-driven colony formation down to the level of control while having minimal impact on control colony formation (Figure 5A, B). Similarly, WM-3835 inhibited *SETBP1^D868N^*-induced colony formation and produced a dose-dependent decrease in colony number (Figure 5D,E). These studies demonstrate selective effects of KAT6/KAT7 inhibitors on SETBP1-mutant cells.

**Figure 5.**
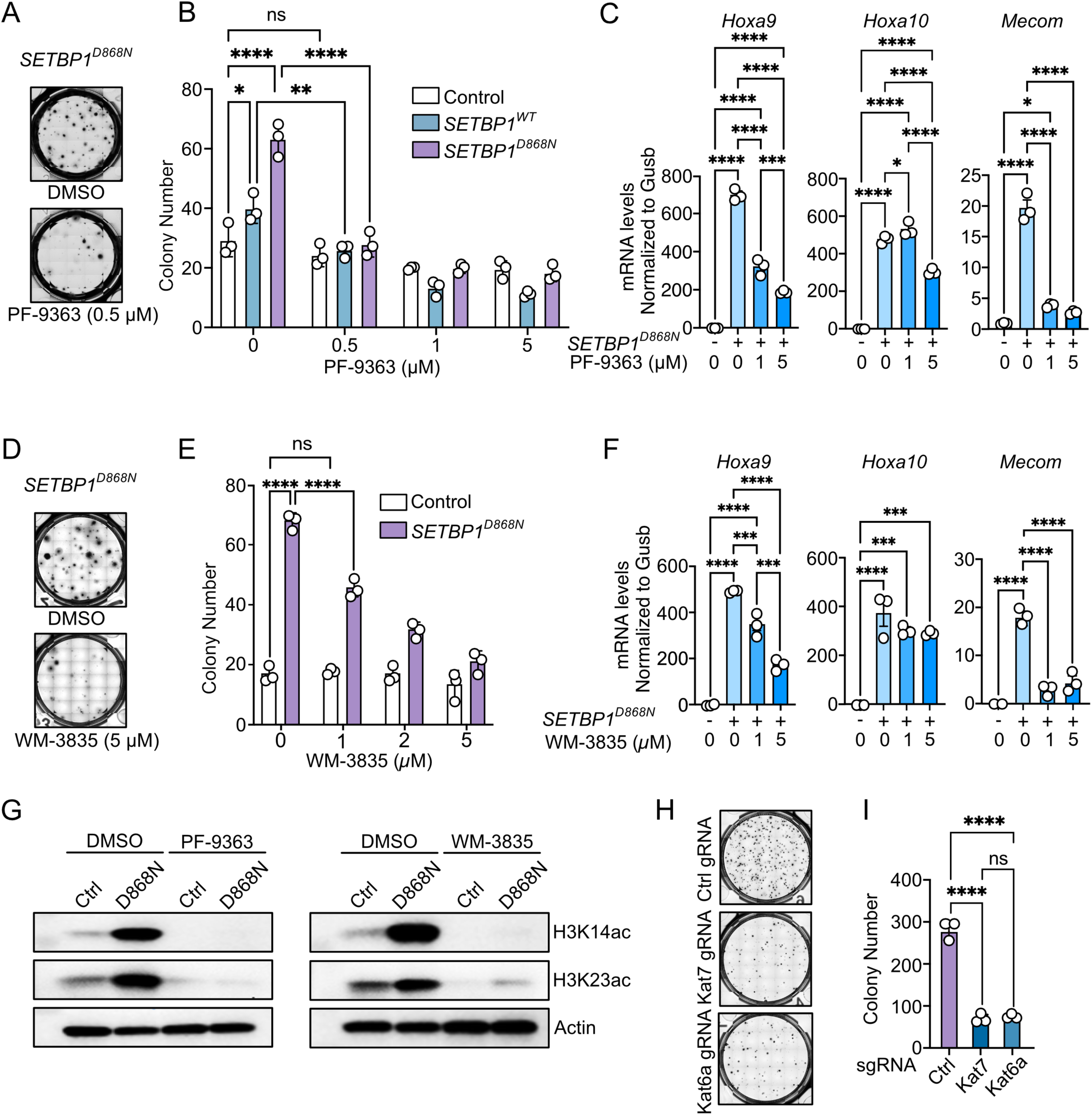
SETBP1-driven cell proliferation is dependent on MYST acyltransferase activity. **A.** Control, *SETBP1^WT^*, *or SETBP1^D868N^*-expressing mouse bone marrow Lin-cells were treated with 0.5, 1, or 5 μM PF-9363 or DMSO (Control) in a colony-forming unit assay. Representative of colonies at day 14. **B.** The number of colonies was counted for each treatment condition. Two-way ANOVA results are shown only for the 0 and 0.5 μM doses for clarity. P-values of higher doses to control had the same level of significance as the lowest dose. **C.** qRT-PCR quantification of *Hoxa9*, *Hoxa10,* and *Mecom* gene levels normalized to *Gusb* control and displayed as fold-change. Mean of 3 biological replicates. **D.** Mouse bone marrow Lin-control or *SETBP1^D868N^*-expressing cells were treated with 1, 2, or 5 μM WM-3835 or DMSO in a colony-forming unit assay. Representative images of 3 biological replicates of colonies at day 14. **E.** The number of colonies was quantified. Two-way ANOVA results shown only for 1 μM dose for clarity. **F.** qRT-PCR quantification of *Hoxa9*, *Hoxa10,* and *Mecom* gene levels normalized to *Gusb* control and represented as fold-change. Mean of 3 biological replicates. **G.** Western blot analysis of H3K14ac and H3K23ac levels in bone marrow Lin-cells treated with PF-9363 (5 μM) or WM-3835 (5 μM) in liquid culture for 5 days. **H.** Lineage-negative HSPCs expressing *SETBP1^D868N^* were sorted, followed by CRISPR/Cas9-based disruption of *Kat7* or *Kat6a*. Representative images of colonies at day 14 are shown. **I.** Colony counts. Error bars represent SEM. Statistics: 1-way ANOVA with Dunnett’s multiple comparisons test. \**p* < 0.05, \*\**p* < 0.01, \*\*\**p* < 0.001, \*\*\*\**p* < 0.0001.

To understand the effects of MYST inhibitors on *SETBP1^D868N^*-driven gene expression programs, RNA was isolated from colonies at 14 days. Levels of *Hoxa9*, *Hoxa10,* and *Mecom* were evaluated by qRT-PCR relative to a housekeeping gene (Gusb) (Figure 5C,F). Both PF-9363 and WM-3835 significantly reduce the expression of these *SETBP1^D868N^* target genes. To confirm the effect of PF-9363 and WM-3835 on histone acetylation in this model, we evaluated the levels of H3K14ac and H3K23ac by immunoblot (Figure 5G). Inhibition of KAT6A and/or KAT7 reversed the increase in these marks driven by *SETBP1*^D868N^ expression. To confirm these findings using a genetic approach, KAT6A and KAT7 were knocked out in SETBP1-mutant cells (Figure S3). Knockdown of either KAT7 or KAT6A resulted in potent inhibition of the colony formation by *SETBP1^D868N^* Lin-hematopoietic progenitors relative to control sgRNAs (Figure 5H, I). Collectively, these studies establish KAT7 and KAT6A as key dependencies in *SETBP1*-mutant cells.

### MYST inhibitors are highly effective against SETBP1-mutant leukemia *in vivo*

To determine whether MYST inhibition is effective against SETBP1-mutant leukemia, we used a *SETBP1^D868N^/CSF3R^T^*^618^*^I^-*driven murine myeloproliferative neoplasm (MPN)^11^. Transplanted mice were treated with PF-9363 (1mg/kg/day) or vehicle by oral gavage for a total of 28 days starting at 8 days post-transplantation (13 mice per group). This *SETBP1*-mutant leukemia model is rapidly evolving. The median survival time for the vehicle-treated cohort was ∼20 days post-transplant (Figure 6A). The PF-9363-treated cohort had a striking improvement in survival, with none of the mice meeting endpoint criteria over the 70-day duration of the experiment. This is remarkable given that the mice only received the drug for 4 weeks.

**Figure 6.**
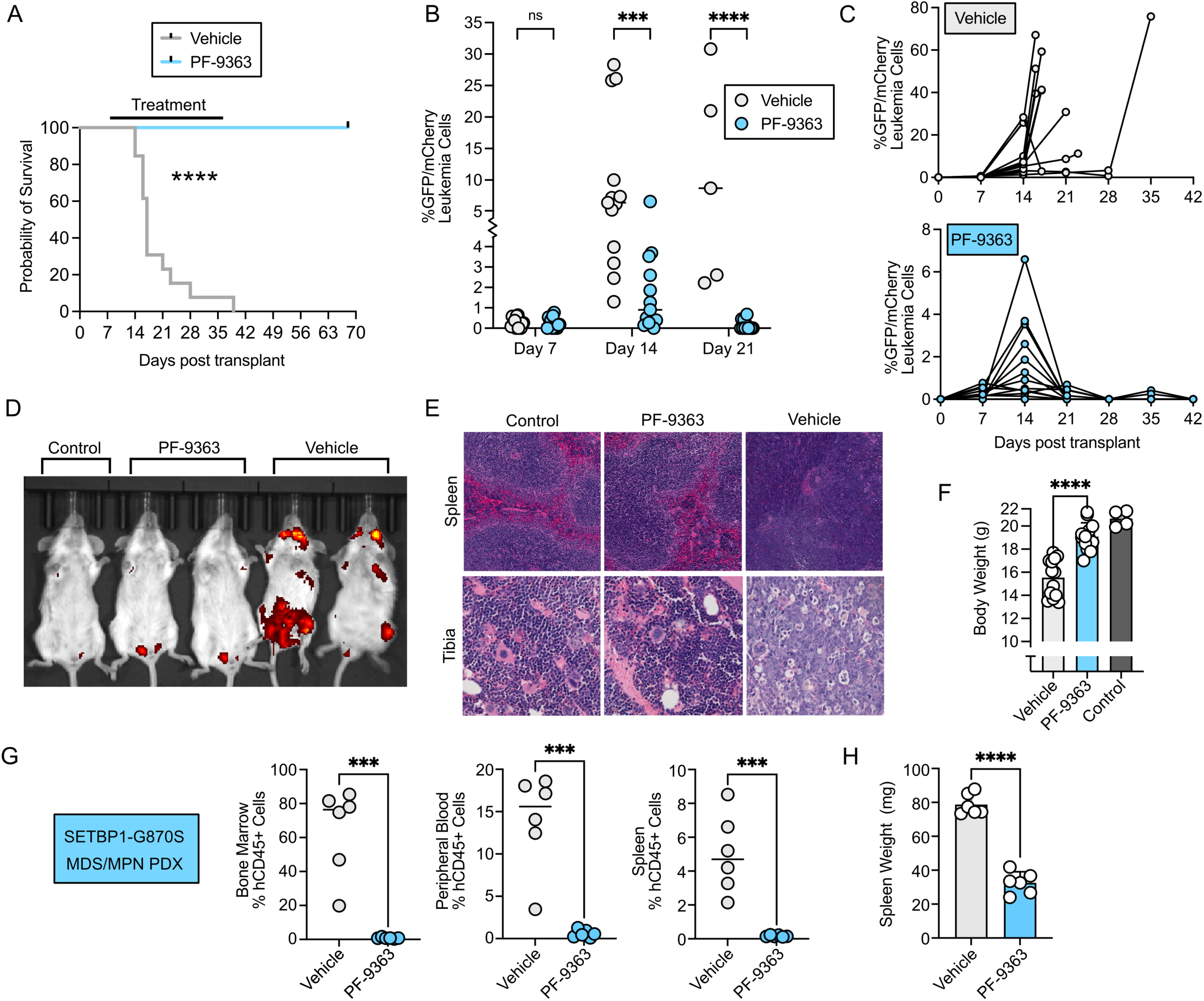
MYST inhibition is effective against *SETBP1*-mutant leukemia *in vivo*. Balb/cJ mice were sub-lethally irradiated and transplanted with 250,000 *CSF3R*^T618I^/*SETBP1*^D868N^ immortalized murine progenitor cells (described previously^11^). These cells have GFP and RFP fluorescent reporters. Mice were treated with Vehicle or PF-9363 (1mg/kg/day) by oral gavage for 28 days starting 8 days post-transplantation. **A.** Kaplan-Meier survival analysis of PF-9363 versus vehicle-treated mice. Statistics: Log-rank (Mantel-Cox) test, \*\*\*\**p* < 0.0001. **B.** Leukemia cell burden was assessed by peripheral blood flow cytometry for GFP+/RFP+ leukemia cells in PF-9363 or vehicle-treated mice (n =13/group) at 7, 14, and 21 days. **C.** Longitudinal tracking of leukemic cell burden by peripheral blood flow cytometry for GFP+/RFP+ cells in individual mice over the course of the treatment window. Please note the differences in scale between the vehicle and PF-9363-treated groups to allow for visualization. **D.** Representative images from the IVIS imager detecting GFP+ cells in mice at 21 days post-transplant and 13 days of drug treatment. Control mice had sub-lethal irradiation, but no leukemia cells were transplanted. **E.** Representative hematoxylin and eosin (H&E)–stained spleen and tibia longitudinal sections. **F.** Body weights of mice at endpoint or day 42 for those mice not reaching endpoint during the study. Control mice were irradiated but did not receive leukemia cells. Statistics: 1-way ANOVA with Dunnett’s multiple comparisons test. \*\*\*\**p* < 0.0001. **G.** Patient-derived xenograft model of an MPN/MDS with a *SETBP1^G^*^870^*^S^* mutation. After confirmation of engraftment of human cells into NSGS mice, mice were treated with PF-9363 at 1mg/kg/day for 4 weeks. The percent of MPN/MDS cells was assessed by flow cytometry with a human CD45 surface marker in the bone marrow, peripheral blood, and spleen. **H.** Spleen weights in the vehicle versus PF-9363 xenografted mice. Statistics: Student’s t-test, \*\*\**p* < 0.001, \*\*\*\**p* < 0.0001.

Engraftment was detectable in both groups at 7 days post-transplant. After just one week of therapy (day 14 post-transplant), mice treated with PF-9363 had a significant decrease in GFP/mCherry-positive leukemia cells relative to the vehicle-treated mice (Figure 6B). This difference was even more pronounced after two weeks of therapy (day 21 post-transplant). White blood cell counts rose in the PF-9363-treated group until day 14, but then sharply declined at day 21 (Figure 6C, bottom panel). In contrast, leukemic cells in the vehicle-treated group continue to rise over time (Figure 6C, top panel). Of note, continued monitoring of the mice by peripheral blood flow cytometry post-treatment did not detect any rebound in GFP/mCherry-positive leukemia levels after cessation of therapy.

Leukemia burden was further evaluated using an IVIS imager after two weeks of therapy (Figure 6D). A strong signal from leukemia cells is observed in the vehicle-treated mice in the long bones, skull, and pelvis. PF-9363-treated mice had markedly reduced leukemia cell burden when compared to the vehicle-treated mice (Figure 6D). In the vehicle-treated cohort, the structure of the splenic white pulp was disrupted, and the clear marginal zones surrounding follicles were hard to visualize, indicating high leukemic burden in the spleen (Figure 6E). In contrast, histological analysis of the spleens of mice that did not receive leukemia cells (control) and the PF-9363-treated cohort revealed a clear distinction between the red and white pulp, with well-defined marginal zones (Figure 6E). Hematoxylin and Eosin staining of the tibias of the vehicle-treated mice revealed hypercellularity, with an abundance of cells with irregularly shaped nuclei with fine chromatin and prominent nucleoli (Figure 6E). In contrast, mice that did not receive leukemia cells, or those treated with PF-9363 treated mice exhibit well-organized bone matrix (stained pink) and cells (nuclei stained blue). The PF-9363-treated mice had significantly higher body weights compared to the vehicle-treated mice, indicating overall better health (Figure 6F). Overall, the PF-9363-treated cohort demonstrated improved physical condition, reduced disease burden, and a striking increase in survival.

To validate the efficacy of MYST inhibition against *SETBP1*-mutant human leukemia, we harnessed patient-derived xenograft models. Bone marrow mononuclear cells from a patient with MDS/MPN with a *SETBP1^G^*^870^*^S^* mutation (functionally analogous to D868N) were engrafted into NSG-S mice. After validation of engraftment by flow cytometry, mice were treated with PF-9363 (1mg/kg) or vehicle control by daily oral gavage for 4 weeks. At the end of the study, the burden of MDS/MPN cells was evaluated by flow cytometry for human CD45. There was a striking reduction in human *SETBP1^G^*^870^*^S^*-mutant leukemia cells in the bone marrow, peripheral blood, and spleen of the PF-9363-treated mice (Figure 6G). Additionally, there was a statistically significant decrease in spleen weight, indicating that the treatment was effective at ameliorating leukemia-associated splenomegaly (Figure 6H). Efficacy was validated in a second CMML PDX model with a *SETBP1^G^*^870^*^S^* mutation (Figure S3D-E). This sample exhibited a significant reduction in human leukemia cells in the spleen and a significant decrease in the percentage of CD34+CD38-CMML stem cells in the marrow following PF-9363 treatment. Collectively, these studies establish MYST acetyltransferases as a therapeutic vulnerability in *SETBP1*-mutant malignancies.

## Discussion

Although *SETBP1* is a myeloid oncogene with strong prognostic significance, a mechanistic understanding of the biology of *SETBP1* mutations has lagged behind other recurrently mutated genes. Induction of stem and progenitor-associated transcriptional programs is a hallmark of *SETBP1*-mutant leukemia (Figure 3)^12,15^, but the precise mechanisms governing these transcriptional changes are incompletely understood. To better understand the mechanisms of transcriptional control, we conducted a screen for SETBP1 interactors (Figure 1). The very high concordance between the SETBP1^WT^ and SETBP1^D868N^ interactomes is consistent with the idea that mutations in SETBP1 serve to increase the levels of SETBP1 available to activate transcription, rather than a neomorphic function.

This screen confirmed a known interaction of SETBP1 with MLL1^15,16^ and, excitingly, identified several novel chromatin regulatory complexes that interact with SETBP1. One of the two topmost enriched interaction partners was BRPF1, a scaffolding protein in MYST acetyltransferase complexes. A subset of MYST complexes contains BRPF1, while others contain alternative scaffolds, such as JADE3. BRPF1 positively regulates Hox gene expression, and its knockout causes severe bone marrow failure^26,27^. There are five MYST acetyltransferases (KAT5, KAT6A, KAT6B, KAT7, and KAT8^28,29^), which have preferences for complexes with particular scaffolding proteins. The proteomics screen revealed significant interactions between SETBP1 and KAT6A and KAT7, two acetyltransferases known to be part of BRPF1-containing complexes^30^. Lacking direct chromatin-binding regions, KAT6A and KAT7 form complexes with BRPF1, EAF6, and ING5 that enable them to interact with target genes^30^. We find that expression of mutant-SETBP1 enhances the co-immunoprecipitation of BRFP1 with KAT7. Since BRFP1 can stimulate MYST acetyltransferase activity^29^, this enhanced interaction could increase the rate of acetylation by these complexes.

Although there may be some overlap in histone mark specificity, evidence suggests that KAT7/BRPF1-containing MYST complexes have a preference for depositing H3K14Ac, while KAT6A/BRPF1 complexes primarily deposit H3K23Ac^24,25,28,30^. Indeed, we find that *SETBP1-mutant* cells have a global increase in deposition of both H3K14Ac and H3K23Ac (Figure 4). This is particularly striking at known SETBP1 target genes, such as the *Hoxa* cluster and *Meis1*. Both H3K14Ac and H3K23Ac function as transcriptional activating marks read by bromodomain proteins. In *SETBP1-mutant* cells, we find that H3K23Ac is almost exclusively promoter localized, while H3K14Ac is present at both promoters as well as enhancers. This increased deposition of activating acetyl marks is consistent with SETBP1 primarily being associated with the induction of target gene expression.

There is a growing interest in the targeting of MYST complexes in cancer. Notably, MYST acetyltransferases positively regulate the expression of the estrogen receptor^31^. A number of MYST inhibitors are now showing promise in clinical trials for breast cancer. In B-cell acute lymphoblastic leukemia, KAT7 promotes leukemogenesis through activation of the Wnt/β-catenin signaling^32^. In hematological malignancies, KAT7 functions as a key regulator in the self-renewal and maintenance of MLL-fusion-transformed leukemia stem cells^33^. Of note, KAT7 and MLL1 directly interact^33,34^, raising the possibility that SETBP1-associated MYST complexes are also linked to MLL1.

Disruption of either the KAT6A or KAT7 genes dramatically reduce SETBP1^D868N^-driven colony formation (Figure 5). Both a putative KAT7 selective inhibitor (WM-3835) and a KAT6A/KAT6B/KAT7 inhibitor (PF-9363) markedly reduce the expression of SETBP1 target genes and suppress SETBP1^D868N^-driven progenitor expansion. Of note, WM-3835 also reduces the KAT6A-deposited histone mark, H3K23Ac, indicating potential for broader activity. PF-9363 had a remarkable efficacy against an aggressive *SETBP1^D868N^* leukemia model. Although this model was uniformly lethal by 6 weeks post-transplant, in the vehicle control group, no mice in the treatment group succumbed to disease, even after discontinuation of therapy. PF-9363 was also highly potent against an MDS/MPN PDX model with a *SETBP1^G^*^870^*^S^*mutation, resulting in the near elimination of malignant cells in the marrow, blood, and spleen.

In this study, we have focused on interactions between SETBP1^D868N^ and MYST complex members. However, our proteomics studies identified a number of transcriptional regulatory complexes that interact with SETBP1. BRD2 and BRD3 were identified as having robust interaction with SETBP1; however, our prior studies found that bromodomain inhibitors failed to normalize SETBP1-associated gene expression programs^11^. This screen also identified an intriguing interaction between SETBP1 and members of the super elongation complex (SEC). MLL is frequently translocated to create a fusion protein involving SEC members^35^. The role of SETBP1 in coordinating MLL and the SEC is an exciting area of future investigation.

Collectively, this study provides us with the first comprehensive map of the SETBP1-interactome and identifies MYST acetyltransferase complexes as a key component of the SETBP1-controlled epigenetic axis. Furthermore, using a combination of in vitro, in vivo, and xenograft models, we identify MYST inhibitors as highly effective against SETBP1-mutant leukemia, thus providing preclinical rationale for the investigation of these drugs in this high-risk patient population.

## Materials & Methods

### Plasmid construction

The human SETBP1^WT^ or SETBP1^D868N^ open reading frame (ORF) with codon optimization were synthesized commercially (Invitrogen/GeneArt). A gateway pENTR vectors were subcloned into the destination tetracycline-inducible lentiviral pCW57.1 (Addgene: 41393) or pSMAL (Addgene 161785) or pcDNA3.1/V5 (ThermoFisher, 12290010) with Gateway cloning (Life Technologies). pSMAL-luciferase was a gift from Peter van Galen. The lentiviral pSTVH7 N-BirA BioID construct was a gift from Anne-Claude Gingras^36^. Gateway-compatible entry clones for the SETBP1^WT^ or SETBP1^D868N^ were fused into the pSTVH7-BirA*-V5 Gateway lentivirus destination vector using Gateway LR Clonase II (Invitrogen). All plasmids were verified by sequencing.

### Cell lines

Doxycycline (DOX) inducible K562 (CCL-243 ATCC) cells were generated by lentivirus transduction. Briefly, 1.0 g of psPAX2 (Addgene #12260), 0.5 g of VSV-G packaging vectors (Addgene #8454) (39), and 1.5 g of pCW57.1-SETBP1^D868N^ were transfected into LentiX cells (Takara) using Fugene 6 reagent as per manufacturer’s recommendations (E2691 Promega). Virus was harvested at 48–72 hours post-transfection, and cells were infected by spinoculation. Transduced cells were selected with puromycin (A1113803 Life Technologies). To generate the BioID cell line, K562 cells were transduced with pSTVH7-BirA* tagged bait (control, SETBP1^WT^ or SETBP1^D868N^) as well as rtTA (Addgene #175274), followed by hygromycin for stable cell line selection. Cell lines were tested for Mycoplasma monthly.

### Antibodies and small molecules

The following antibodies were used in this study: anti-SETBP1 (16841-1-AP Proteintech); anti-KAT7(58418 Cell Signaling Technology (CST)); anti-H3K14 ac (7627 CST); anti-H3K23ac (86664 CST); anti-KAT6A (78462 CST), anti-DOT1L (90878 CST); anti-KMT2A (14197 CST); anti-KMT2B(47097 CST); anti-IgG (2729 CST); anti-rabbit IgG, HRP-linked antibody(7074 CST); anti-BRPF1(259840 Abcam); Anti-V5 antibody (EPR278187-61 Abcam).

The small-molecule inhibitors used in this study were purchased from MedChemExpress. They are WM-3835 (HY-134901) and PF-9363 (HY-132283). All other reagents were of analytical grade from Sigma.

### Proximity-dependent biotin identification (BioID)-based sample preparation

BioID methodology performed as described by Sears et al^37^. Briefly, BioID control, *SETBP1^WT^*, and *SETBP1^D868N^*K562 cell lines were selectively grown in the presence of 200μg/mL hygromycin up to 80% confluence before expression was induced via 1μg/mL tetracycline and 50μM biotin for 24 hours. Cells were pelleted at low speed and washed with ice-cold PBS. Cell pellets were lysed with 8M urea containing protease inhibitor cocktail (P8340 Sigma-Aldrich, 1:500) and Pierce Universal Nuclease for 10 minutes at 25 °C. 20% Triton X-100 was added to cell lysates, then samples were sonicated at 4 °C using three 10-second bursts with 10-second pauses at 30% amplitude with a QSonic. The lysate was cleared by centrifugation at 16,500 x g for 10 minutes at 4 °C. Samples were preincubated with gelatin Sepharose beads for 2 hours at 4 °C and then with prewashed streptavidin Sepharose beads for 4 hours at 4 °C on a rotator. After incubation, beads were washed three times with 8 M urea in 50 mM Tris-HCl buffer (pH 8.0), followed by three washes with 50 mM ammonium bicarbonate (ABC) (pH 8.5). Proteins bound to beads were digested in 200 μL of 50 mM ABC containing 1 μg of trypsin (V5280 Promega) at 37 °C overnight with gentle agitation. A second digestion step was performed by adding 0.5 μg of trypsin in 10 μL of 50 mM ABC and incubating for an additional 2 hours at 37 °C. After digestion, the supernatant was collected, and the beads were rinsed twice with 150 μL of mass spectrometry-grade water. The rinses were combined with the initial digest eluate, and the pooled samples were stored at -80 °C until ready for mass spectrometry analysis. Mass spectrometry data acquisition and processing are described in detail in the supplemental methods.

### BioID validation, co-immunoprecipitation, and western blot

To validate the BioID results, K562 cells transduced with pSTVH7-BirA*-SETBP1-BirA were induced using 1 μg/ml DOX, harvested, and lysed with ice-cold Pierce RIPA buffer (PI89900PM) plus protease inhibitor cocktail (P8340 Sigma-Aldrich). The samples were sonicated at 4 °C as described above. The lysates were centrifuged at 16,000 g for 20 minutes at 4 °C, and the supernatant added to tubes containing 60 μL of pre-washed streptavidin-sepharose bead slurry. Samples were rotated for 3 hours at 4 °C. The beads were gently pelleted and washed 4 x 1 mL with RIPA buffer. Following the final wash, the beads were eluted with 45 μl 1X GS buffer (2% sodium dodecyl sulfate, 50 mM Tris pH8, 10% glycerol, and 0.03% bromophenol blue) plus 5% 2-mercaptoethanol (21985023 ThermoFisher) and prepared for SDS-PAGE.

For co-immunoprecipitation studies, pcDNA3.1/V5-control or pcDNA3.1/V5-SETBP1^D868N^ plasmids were transfected into HEK293 cells with Fugene 6. Cells were harvested 48 hours with Pierce IP lysis buffer (PI87788). 1mg of total cell lysate was incubated with anti-V5 antibody overnight at 4 °C. The lysates were incubated with prewashed protein A/G plus-agarose (sc-2003 Santa Cruz Biotechnology) for 3 hours at 4°C. Beads were washed with cold IP buffer 4 times to remove unbound proteins and centrifuged at 2500 rpm for 5 minutes at 4 °C. The samples were eluted and prepared for SDS-PAGE. Scientific SuperSignal West Pico PLUS Chemiluminescent (34580 ThermoFisher) was used for detection.

### Murine hematopoietic colony-forming unit (CFU) assays

C57BL/6J mice (#000664) were obtained from The Jackson Laboratories. Female mice were used between 8-9 weeks of age. All murine experiments were conducted in accordance with the National Institutes of Health Guide for the Care and Use of Laboratory Animals and approved by the Institutional Animal Care and Use Committee of Oregon Health & Science University (Protocol #TR03_IP00000482).

CFU assays were performed as described previously^38^. Lentivirus were generated by co-transfecting psPAX2 (Addgene: 12260), pMD2.G (Addgene: 12259), and pSMAL_Luciferase (control) or pSMAL_SETBP1^D868N^ into LentiX cells (Takara) with Fugene 6 (Promega E2691), according to the manufacturer’s instructions. Viral supernatants were harvested 48 and 72 hours later. Murine bone marrow cells were isolated from 8 weeks old female C57BL/6J mice. Lineage negative cells (Lin-) were isolated with Direct Lineage Cell Depletion Kit (130-110-470 Milternyi Biotech) following manufacture’s protocol, were cultured in Prestim medium (IMDM medium with 15% FBS presence of SCF, 50ug/ml IL6, and IL3). Lin-cells were spinnoculated with viral supernatant on two subsequent days (2,500 rpm for 90 minutes at 30 °C, brake turned off). Positive GFP cells were sorted on a Sony sorter into MethoCult M3534 methylcellulose (StemCell Technologies). 1000 Lin-cells were used per replicate. Colonies were imaged using STEMvision (StemCell Technologies), blinded, and then manually counted using ImageJ (NIH).

### *SETBP1*-mutant murine leukemia transplantation and drug treatment

8-week-old female Balb/cJ mice (#000651 The Jackson Laboratories) were sub-lethally irradiated (5Gy) and 250000 *CSF3R^T^*^618^*^I^*/*SETBP1^D868N^* murine bone marrow progenitor-derived cells^8^ (GFP and mCherry positive) were retro-orbitally injected. After transplantation, mice were maintained on antibiotic water for two weeks (Polymyxin B sulfate salt, P1004; NeoMycin trisulfate salt hydrate, N1876). Experimental animals were randomized into treatment groups and dosed once a day by oral gavage with 1mg/kg PF9363 or vehicle (5% DMSO/40% PEG300/55% 1 x PBS). As a toxicity control, 4 mice were irradiated, receiving no cell transplantation, and gavaged with PF-9363. Drug treatment started 8 days after transplant and continued for 4 weeks. Peripheral blood collected from the saphenous vein was monitored weekly for white blood cell counts (WBC) using a Vet ABC animal blood counter (Scil animal care company). Flow analysis was performed weekly and at the time of humane endpoint, to assess the percentage of GFP/mCherry positive cells on an LSRFortessa analyzer (BD). For imaging, randomized mice from each group were placed under isoflurane-induced anesthesia and oriented ventral side up to facilitate oxygen and isoflurane flow through the nose cones. Images were taken using IVIS Spectrum II in *vivo* imaging system and analyzed with Living Image 4.2 Software (Revvity).

### Patient-derived xenografts

Patient bone marrow samples were obtained with informed consent in accordance with the Moffitt Cancer Center and the University of South Florida Institutional Review Boards in accordance with the Declaration of Helsinki. Xenotransplantation was performed as described previously^39^. Mononuclear cells were isolated using Ficoll-Paque followed by RBC lysis from an MPN/MDS and CMML bone marrow aspirates. The mutational profile for MDS/MPN includes the following mutations: SETBP1^G870S^, ASXL1^G646Wfs*12^, U2AF1^R156H^, KDM6A^R1213*^, ETV6^R399C^, and FLT3^D835E/N676K^. Mutations detected in the CMML-0 sample were *SETBP1^G^*^870^*^S^*, *SRSF2^P95H^*, *ASXL1^Y^*^974^*\**, *TET2^F^*^1285d^*^el^*, and *FLT3^N^*^676^*^K^*. Female, 7-week-old, NOD-scid IL2Rgnull-3/GM/SF (NSG-S) mice were irradiated (250 rads) the day before tail vein injection of 4.5×10^6^ cells. Flow cytometry for human CD45 on peripheral blood was used to confirm engraftment. Mice were then randomized to vehicle and treatment groups and treated with 1mg/kg/day PF-9363 for 4 weeks as described above. After four weeks of treatment, leukemic disease burden in the spleen, peripheral blood, and bone marrow was assessed by flow cytometry.

### Flow cytometry for xenograft models

Staining was performed in 5 mL polystyrene test tubes with 2 washes per wash step. Red blood cells in bone marrow, spleen, and peripheral blood samples were lysed by incubating the samples for 5 minutes in Ammonium Chloride Potassium (ACK) lysis buffer. Cells were then washed and stained with LIVE DEAD violet fixable dye (Thermo Fisher, Cat# L34964) for thirty minutes, after which they were fixed with 1.6% formaldehyde for ten minutes at room temperature. Following this, cells were washed and incubated with mouse and human FcR blocking reagents (Miltenyi Biotec Cat# 130-092-575 and 130-059-901) for ten minutes then incubated with fluorophore-conjugated antibodies for thirty minutes in BD brilliant Stain Buffer (BD Biosciences Cat# 566349) and washed. Antibodies used are as follows: PE mouse anti-human CD34 (BD Biosciences Cat# 555822), BB515 mouse anti-human CD33 (BD Biosciences Cat# 564588), PerCP-Cy5.5 mouse anti-human CD19 (BioLegend Cat# 363016), BV421 anti-mouseTer-119 (BioLegend Cat# 116234), BV605 mouse anti-human CD45 (BD Biosciences Cat# 564047), BV711 mouse anti-human CD16 (BD Biosciences Cat# 563127), APC mouse anti-human CD3 (BD Biosciences Cat# 555342), PE 594 mouse anti-huma CD38 (BioLegend Cat# 303538), BUV395 mouse anti-human CD14 (BD Biosciences Cat# 563561), BUV737 mouse anti-mouse CD45.1 (BD Biosciences Cat# 612811).

### Histology and microscopy

Tibias and spleens were fixed with 10% formalin for 24 hours. After 24 hours, spleens were transferred to 70% ethanol until processing. Tibias were decalcified using 10% EDTA for 5 day and then stored in 70% ethanol. Tibias and spleens were then embedded in paraffin, sectioned, and stained using H&E. Images were taken on an ApoTome3 - Zeiss Microscope with Grid-Based Optical Sectioning.

### RNA isolation, qRT-PCR, and bulk RNA-sequencing

Briefly, cells from CFU were isolated from methylcellulose and washed well with PBS. Total cellular RNA was isolated with RNeasy mini kit (74106 Qiagen) according to the manufacturer’s instructions. 500ng of total RNA was used for first-strand cDNA using a High-Capacity cDNA reverse transcription kit (4368814 ThermoFisher) following the manufacture’s protocol. qPCR was performed on QuantStudio 7 Flex Real-Time PCR System (Applied Biosystem) using taqman probes (ThermoFisher) for Hoxa9 (Mm0043964), Hoxa10 (Mm07295769), and Mecom (Mm00491303) with TaqMan™ Universal PCR Master Mix (4364349 ThermoFisher).

For bulk RNA-seq, 5 x10^5^ lineage-depleted bone marrow cells were transduced with pSMAL_luciferase or pSMAL_SETBP1^D868N^ and sorted for GFP-positivity (as described above). RNA was extracted as described above. mRNA libraries were prepared using a Watchmaker mRNA Library Prep Kit (7K0105-024) per the manufacturer’s protocol. 500ng RNA for each sample was used for mRNA capture. xGen Stubby Adapter was used in ligation, and UDI primers were used for library amplification (Integrated DNA Technology # 10005976). Libraries were sequenced on a NextSeq 2000 (Illumina) using the NextSeq 1000/2000 P2 kit with paired-end 2 x 61 bp reads. Analysis is described in the supplemental methods.

### CUT&RUN

CUT&RUN was performed with Epicypher CUT&RUN kit. Briefly, 500,000 cells immobilized with Concanavalin A magnetic beads were permeabilized with Digitonin (0.01% for K562 cells and 0.002% for mouse Lin-cells) and incubated overnight at 4 °C with 0.5 μg antibody. After the addition and activation of pAG-MNase, CUT&RUN-enriched DNA was purified using SPRIselect beads. 5ng DNA was used to prepare sequencing libraries with NEBNext Ultra II DNA Library prep (New England Biolabs E7645). PCR conditions were set at 98 °C for 30 sec initial denaturation, 13X 98 °C for 10 sec and 65°C for 10 sec, and then 65 °C for 5 minutes for final extension. Libraries were analyzed by Agilent TapeStation, pooled to equivalence, and sequenced on NextSeq 500, NextSeq 2000, or NovaSeq 6000 sequencers (Illumina). Analysis is described in the supplemental methods.

### CRISPR/Cas9

For CRISPR-based gene knockout in mouse Lin-cells EditCo Gene Knockout Kits were used. Kat7 and Kat6a triple-sgRNA sequences are listed in Supplementary Table 1. SETBP1^D868N^-transduced and GFP-sorted Lin-cells were used in this experiment. Briefly, 30μM sgRNA was combined with Cas9 (9:1 ratio) in 20 µl P3 primary buffer (V4XP-3032 Lonza) and allowed to form ribonucleoprotein (RNP) complexes for 30 min. To the RNP solution, 5 µL of cells resuspended at 5 × 10^4^ cells/µL was added and gently mixed. Cells were then electroporated using a 4D-Nucleofector (Lonza) with program CM-137. Following transfection, cells were recovered in culture media and plated into 24-well plates. After 72 hours, replicate samples were either plated CFU assay (5000cells/well), or harvested to analyze knockout efficiency. Genomic DNA was isolated using QuickExtract (Lucigen) following the manufacturer’s protocol. An amplicon containing the target sequence was amplified via PCR with AmpliTaq Gold 360 Master Mix (ThermoFisher) using primer sequences listed in Supplementary Table 1. PCR amplicons were analyzed by Sanger sequencing (Genewiz), and the resulting traces were deconvolved with Synthego’s Inference of CRISPR Edits (ICE) program (https://ice.synthego.com).

## Acknowledgements

Funding for this work was provided by a Blood Cancer United Scholar Award, a Vera and Joseph Dresner Foundation MDS Research Fund Established Investigator Award, The Andrew McDonough B+ Foundation Childhood Cancer Research Grant, a Kuni Foundation Imagination Award, the Knight Cancer Institute NCI Cancer Center Support Grant P30CA069533, and generous donations made possible through the OHSU Knight Cancer Institute to JEM. ST is supported by a Blood Cancer United Fellow Award. SAC was supported by NRSA F32 CA239422. The authors thank the following Oregon Health and Science University core facilities for their assistance: OHSU Flow Cytometry and Monoclonal Antibody Shared Resource (RRID:SCR_009974), Advanced Light Microscopy (RRID:SCR_009961), Histopathology Shared Resource (P30 CA069533 and P30 CA06953313S5). Short read sequencing assays were performed by the OHSU Massively Parallel Sequencing Shared Resource. We gratefully acknowledge Andrew Adey’s lab at OHSU for providing access to their sequencing equipment, which was used for one of the datasets in this study. The research reported in this publication used computational infrastructure supported by the Office of Research Infrastructure Programs, Office of the Director, of the National Institutes of Health under Award Number S10OD034224. The content is solely the responsibility of the authors and does not necessarily represent the official views of the National Institutes of Health. PDX models were collected under IRB protocol MCC 22357 run by Dr. David Sallman at Moffitt Cancer Center. Portions of the experimental work were supported by the Environmental Molecular Sciences Laboratory (grid.436923.9), a U.S. Department of Energy (DOE) Science User Facility at the Pacific Northwest National Laboratory (PNNL), sponsored by the Office of Biological and Environmental Research. PNNL is a multiprogram national laboratory operated by Battelle for the DOE under contract DE-AC05-76RL01830.

## Author notes

The sequencing data generated in this study have been deposited in the Gene Expression Omnibus (GEO) under accession numbers GSE303625 and GSE303627.

---

**Supplemental Figure 1.**
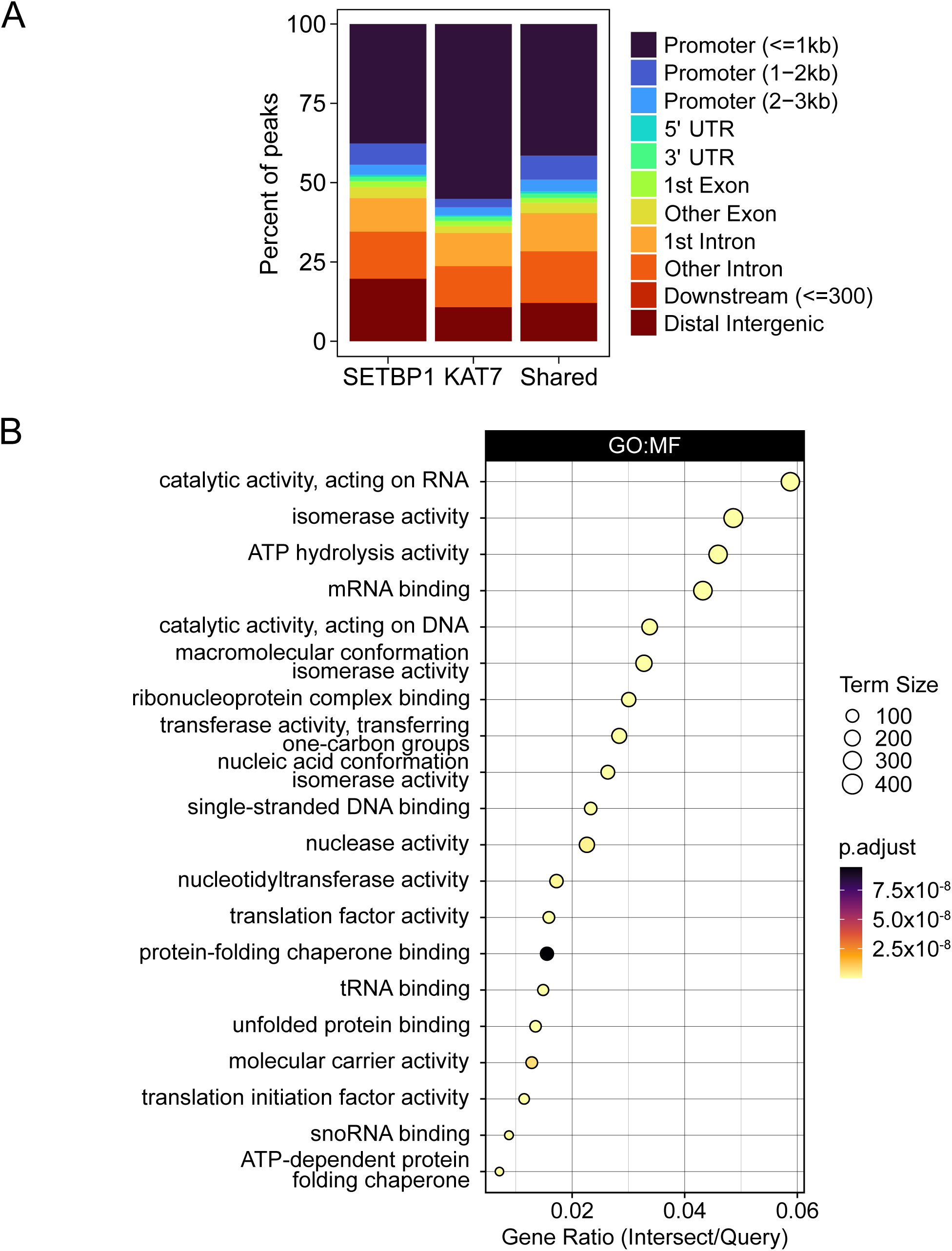
**A.** Genomic feature distribution for peaks in SETBP1, KAT7, and shared (co-occupied) regions in SETBP1^D868N^-induced condition (K562 cells). Approximately 50% of the shared regions overlap with promoter regions. **B.** Top 20 Gene Ontology (GO) Molecular Function (MF) terms from over-representation analysis of SETBP1^D868N^ upregulated genes (padj<0.05) based on RNAseq data. GO terms were selected based on adjusted p-value and ordered by gene ratio for visualization. Gene ratio indicates the proportion of input upregulated genes in each GO term (intersect) relative to the total input upregulated genes (query). Term size represents the total number of genes annotated to each GO term. Color reflects the adjusted p-value of functional enrichment.

**Supplemental Figure 2.**
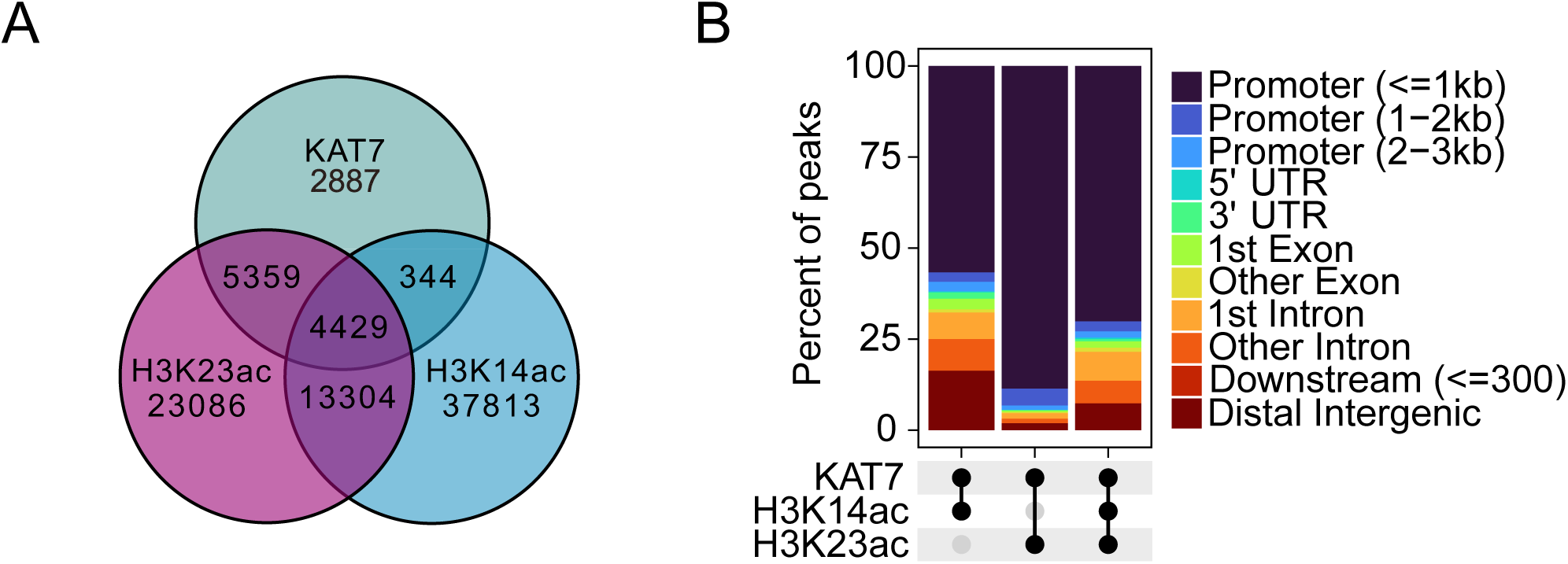
**A.** Supplemental analysis of CUT&RUN data from control or *SETBP1^D868N^*-expressing murine progenitors. **A.** Venn diagram illustrates overlapping regions for KAT7, H3K14ac, and H3K23ac peaks in Lin-cells transduced with *SETBP1^D868N^*. **B.** Genomic feature distribution for KAT7-shared (co-occupied) regions in the SETBP1^D868N^ condition. The leftmost bar corresponds to regions that are co-occupied by KAT7 and H3K14ac, but not H3K23ac. The middle bar corresponds to regions that are co-occupied by KAT7 and H3K23ac, but not H3K14ac. The rightmost bar corresponds to regions that are co-occupied by KAT7, H3K14ac, and H3K23ac.

**Supplemental Figure 3.**
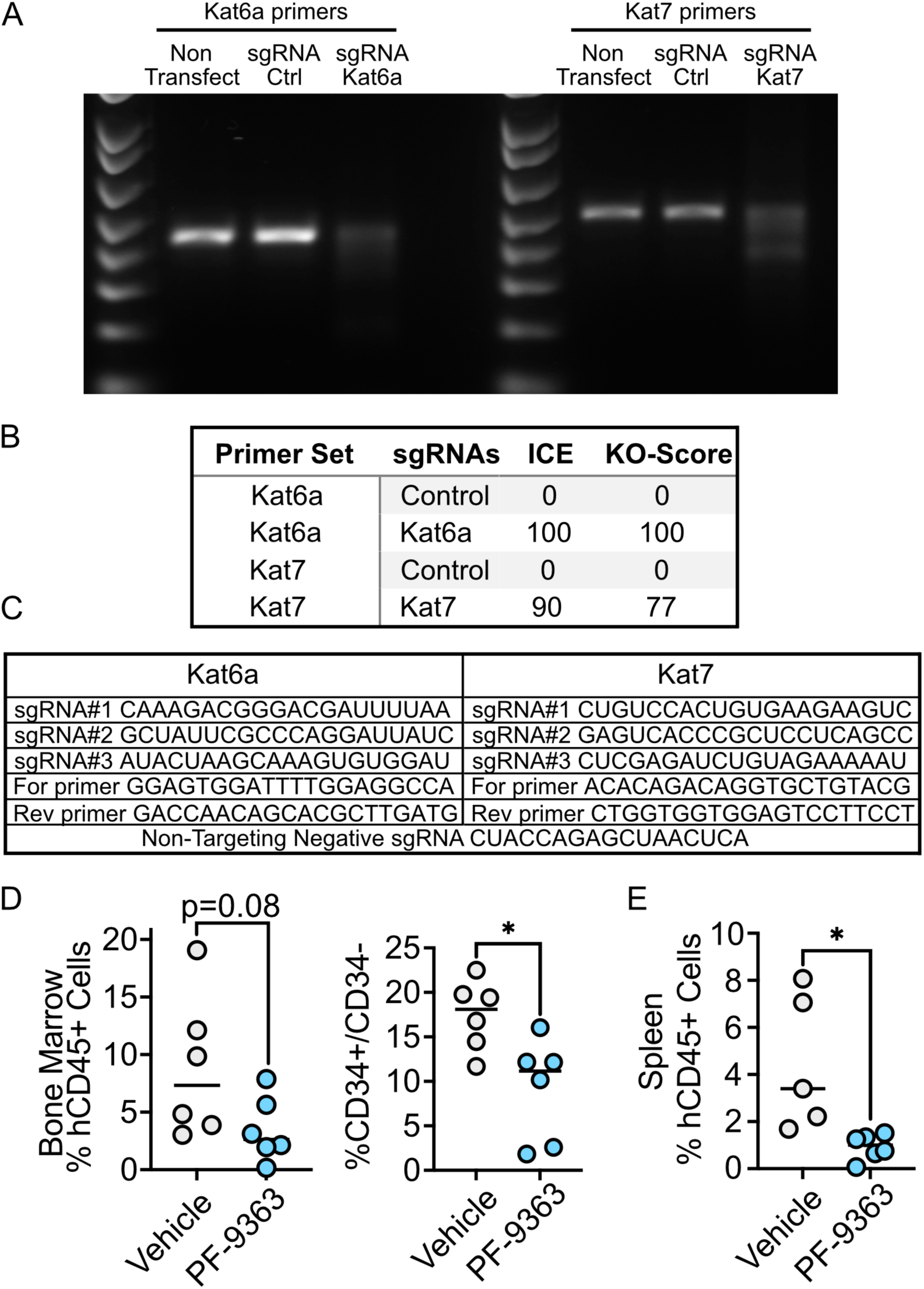
Validation of CRISPR/Cas9-mediated genome editing. The Cas9 nuclease was targeted to Kat6a or Kat7 genes by selected triple-sgRNAs in SETBP1^D868N^ Lin-cells. A. PCR products amplified from non-transfected, electroporated sgRNA control, or sgRNA Kat6a and sgKat7 cells. B. Inference of CRISPR Editing (ICE) and Knockout (KO) score for Sanger sequenced PCR products. C. sgRNA and primer sequences. D. PDX xenograft model from a CMML sample with a *SETBP1^G^*^870^*^S^* mutation along with *SRSF2^P95H^*, *ASXL1^Y^*^974^*\**, *TET2^F^*^1285d^*^el^*, and *FLT3^N^*^676^*^K^*. Flow cytometry-based analysis of the engraftment of human cells into the bone marrow in the control and PF-9363-treated groups, with a reduction in the CD34+/CD38-derived hematopoietic stem cells. Reduced human xenograft cells in the spleen by flow cytometry. Of note, this PDX sample did not cause splenomegaly, likely because the cells have a different tropism than the PDX model in Figure 6.

## SUPPLEMENTAL METHODS

### RNA-seq analysis

Reads were trimmed using Fastp^1^ with options ‘--detect_adapter_for_pe’, ‘--trim_poly_g’, and ‘--adapter_fasta’ using a custom FASTA file of adapter sequences provided by the library kit manufacturer. Trimmed reads were aligned to the mouse genome (GRCm38/mm10) using STAR^2^. A raw gene-level counts matrix was generated from STAR-aligned reads, and DESeq2^3^ was used for normalization and differential expression analysis. Transcription factor activity was predicted from normalized RNA-seq counts using Priori^4^. Gene Set Enrichment Analysis (GSEA)^5,6^, was performed using mouse hallmark (MH) and curated (M2) gene sets from the Molecular Signatures Database (MSigDB)^7^.

Over-representation analysis of SETBP1^D868N^-upregulated genes (padj < 0.05) was performed using g:Profiler, and Gene Ontology (GO)^8,9^ term enrichment results (padj < 0.05) were used for interpretation. GO terms with fewer than 15 genes or more than 500 genes were filtered from g:Profiler results to remove noise from overly narrow or broad categories. Semantic similarity analysis of remaining g:Profiler results was performed with GOSemSim^10,11^, using Wang’s graph-based measurement algorithm^12^. To reduce redundancy, all semantically similar GO terms (similarity > 0.7) were grouped together, and the term with the most significant p-value in each group was retained. From this curated list of GO terms, those with the most significant p-values were selected for visualization.

### CUT&RUN analysis

Reads were trimmed using Fastp^1^ with options “--detect_adapter_for_pe”, “--trim_poly_g”, and “--adapter_fasta” using a custom FASTA file of adapter sequences provided by the library kit manufacturer. Trimmed reads from K562 samples were aligned to the human genome (GRCh38/hg38) and trimmed reads from murine Lin-samples were aligned to the mouse genome (GRCm38/mm10) using Bowtie2^13^ with parameters “--local --very-sensitive-local --no-unal --no-mixed --no-discordant --phred33 -I 10 -X 700”. Duplicates were marked using the Sambamba^14^ “markdup” subcommand. Peaks were called using GoPeaks^15^ using IgG samples as negative controls. To define consensus regions for each antibody target, peaks present in at least two replicates per sample condition were retained as high-confidence regions. These reproducible peaks were then merged across conditions using BEDTools^16^ to generate a unified consensus peak set for each target.

Signal at consensus regions was quantified and analyzed using BEDTools and deepTools^17^. To generate counts-per-million (CPM) normalized bigWig tracks from BAM files, we used deepTools bamCoverage with the parameters “--binSize 10 --smoothLength 50 --normalizeUsing CPM”. Coverage tracks were then averaged across replicates within each sample group using WiggleTools^18^ and the UCSC bedGraphToBigWig tool^19^. To generate heatmaps and median signal profiles of normalized CUT&RUN signal in regions spanning +/- 3kb from consensus region centers, we used deepTools computeMatrix with parameters “--referencePoint center --upstream 3000 --downstream 3000” and deepTools plotHeatmap with parameter “--averageTypeSummaryPlot median”.

For K562 samples, motif analysis of high-confidence peaks in the SETBP1^D868N^-induced condition was performed using HOMER^20^. To assess global similarity across antibody targets (KAT7, SETBP1, KMT2A), a union peak set was generated with BEDTools by merging reproducible peaks (peaks present in at least two replicates) from the SETBP1^D868N^-induced (+DOX) condition for each target. Read counts across this union peak set were quantified using DiffBind with normalization by reads in peaks^21^. For each target, normalized counts from replicates were averaged to create a representative profile. Spearman rank correlation was computed on these averaged profiles using R statistical software. The resulting correlation matrix was visualized with hierarchical clustering using ComplexHeatmap^22^.

To identify co-bound (overlapping) regions between specific antibody targets, reproducible peaks from the +DOX condition were used for K562 samples, and reproducible peaks from the SETBP1^D868N^ condition were used for mouse Lin-samples. For samples from each cell type, overlapping regions between antibody targets were identified using the BEDTools “multiinter” subcommand. The resulting intersection files were used as input to Intervene^23^ to generate Venn diagrams visualizing shared binding regions.

ChIPseeker^24^ was used with the Bioconductor annotation packages “TxDb.Hsapiens.UCSC.hg38.knownGene” (human) or “TxDb.Mmusculus.UCSC.mm10.knownGene” (mouse) to annotate the nearest genomic features and calculate the genomic distribution of reproducible peaks and co-bound regions of interest. For K562 samples, regions from the +DOX condition were used, and for mouse Lin-samples, regions from the SETBP1^D868N^ condition were used.

Replicate-averaged, CPM-normalized bigWig tracks were visualized at loci of interest using the Integrative Genomics Viewer (IGV) web application^25^.

### Mass spectrometry data acquisition

Peptides were resuspended with in 10 μL of 3% acetonitrile (ACN) with 0.1% formic acid (FA), and 3 μL were injected for analysis using a nanoAquity UPLC system (Waters Corporation) coupled to an Orbitrap Fusion Lumos mass spectrometer (Thermo Fisher Scientific). Peptide separation was performed on an in-house packed analytical column (75 μm i.d. x 200 mm) using ReproSil-Pur 120 C18-AQ 1.9-μm resin (Dr. Masch GmbH, Germany). A 120-minute reverse-phase gradient was applied at a flow rate of 200 nL/min with mobile phase A (0.1% FA in water) and mobile phase B (0.1% FA in ACN). The gradient included a linear increase from 2% to 30% B over 84 minutes, followed by a ramp to 90% B and re-equilibration to initial conditions.

Mass spectrometry (MS) was conducted in data-dependent acquisition (DDA) mode. The electrospray voltage was maintained at 2.2 kV with the ion transfer tube temperature at 250 °C. Full MS scans were acquired over an m/z range of 350–1800 at a resolution of 60,000 with an automatic gain control (AGC) target of 4 × 10⁵ and a maximum injection time of 50 ms. MS/MS spectra were acquired for selected precursors using quadrupole isolation (0.7 m/z), higher-energy collisional dissociation (HCD) at 30% normalized collision energy, and Orbitrap detection at 50,000 resolution with an AGC target of 1 × 10⁵ and a maximum injection time of 105 ms. The total duty cycle was set to 2 s.

### Mass spectrometry data processing

Raw MS data were processed using MaxQuant (ver. 1.6.12.0)^26,27^ with the Andromeda search engine, searching against the Human Uniprot protein sequence database (April 12, 2017 with 20,198 sequences). Default search parameters were used, including precursor mass tolerances of 20 ppm for the first search and 4.5 ppm for the main search. The enzyme was set to trypsin, allowing up to two missed cleavages. Carbamidomethylation of cysteine was specified as a fixed modification, and oxidation of methionine and N-terminal acetylation were set as variable modifications. Peptide-spectrum matches (PSMs), peptides and protein groups were sequentially filtered at a false discovery rate (FDR) of 1%. The resulting Protein groups file was analyzed using Perseus software^28^. Proteins identified by only a modification site, reverse decoy sequences, or potential contaminants were excluded. Protein intensities were log2 transformed, and proteins quantified in at least three samples within one group were filtered. Missing values were imputed based on a normal distribution with a downshift of 1.8 standard deviations and a width of 0.3 relative to each sample’s distribution. Two-sided t-tests were performed for pairwise comparisons, and significant proteins were determined using a q-value threshold of 0.05. The MS proteomics raw data have been deposited to the MassIVE (Mass Spectrometry Interactive Virtual Environment) open access repository under the data set identifier MSV000098305 (https://massive.ucsd.edu/ProteoSAFe/static/massive.jsp).

